# Social network analysis and the implications for Pontocaspian biodiversity conservation in Romania and Ukraine: A comparative study

**DOI:** 10.1101/740084

**Authors:** Aleksandre Gogaladze, Niels Raes, Koos Biesmeijer, Camelia Ionescu, Bianca Pavel, Mikhail O. Son, Natalia Gozak, Vitaliy Anistratenko, Frank P. Wesselingh

## Abstract

Romania and Ukraine share the Black Sea coastline, the Danube Delta and associated habitats, which harbor the unique Pontocaspian biodiversity. Pontocaspian biota represents endemic aquatic taxa adapted to the brackish (anomalohaline) conditions, which evolved in the Caspian and Black Sea basins. Currently, this biota is diminishing both in the numbers of species and their abundance because of human activities. Consequently, its future persistence strongly depends on the adequacy of conservation measures. Romania and Ukraine have a common responsibility to effectively address the conservation of this biota. The socio-political and legal conservation frameworks, however, differ in the two countries - Romania is a member of the European Union (EU), thus complying with the EU environmental policy, whereas Ukraine is an EU-associated country. This may result in differences in the social network structure of stakeholder institutions with different implications for Pontocaspian biodiversity conservation. Here, we study the structure and implications of the social network of stakeholder organizations involved in conservation of Pontocaspian biodiversity in Romania, and compare it to Ukraine. We apply a mix of qualitative and quantitative social network analysis methods to combine the content and context of the interactions with relational measures. We show that the social networks of stakeholder organizations in Romania and Ukraine are very different. Structurally, in Romanian network there is a room for improvement through e.g. more involvement of governmental and non-governmental organizations and increased motivation of central stakeholders to initiate conservation action, whereas Ukrainian network is close to optimal. Regardless, both networks translate into sub-optimal conservation action and the road to optimal conservation is different. We end with sketching implications and recommendations for improved national and cross-border conservation efforts.

## Introduction

Pontocaspian (PC) biota is a unique, endemic flora and fauna which includes mollusks, crustaceans, planktonic groups (e.g. dinoflagellates and diatoms) and fish species. This biodiversity evolved in brackish (anomalohaline) conditions of the Black Sea and Caspian Sea basins over the past 2.5 million years [1,2] and nowadays PC communities inhabit the Northern Black Sea, Sea of Azov, Caspian Sea and adjacent river and lake systems, stretching across the vast political and administrative boundaries of the surrounding countries [3]. Currently, PC biota is decreasing in numbers of species and abundances as a result of human activities and their future persistence strongly depends on the adequacy of conservation measures [1,4,5]. Romania and Ukraine hold an important part of the PC habitats. PC species in Romania are limited to the Razim-Sinoe-Babadag lake complex [6,7], the area along the Danube River and the Black Sea coastal zone, which together form the Danube Delta and have the status of Biosphere Reserve. In Ukraine, PC communities occur in the coastal lakes, deltas and estuaries from the Danube Delta to the Dnieper estuary as well as estuarine/coastal habitats in the northern Sea of Azov [8-10]. The two countries share the responsibility of conserving the shared PC habitats and the associated threatened biota [7,9,11,12], however, they have very different socio-political settings and cultural background - Romania became member of the European Union (EU) in 2007, thus complying with the EU environmental policy, and Ukraine is an association agreement signatory non-EU country since 2017.

In both countries Pontocaspian species are threatened and the conservation measures are urgently required. In the past 30 years, the number, abundance and distribution ranges of PC species have decreased dramatically in Romania as a result of human influence [6,7]. In Ukraine, PC species are declining as a result of habitat fragmentation caused by dam and deep sea shipping lane constructions [13,14]. Some of the PC species (e.g. some mollusk and sturgeon species) are of national concerns in both countries - they are recognized to be threatened and in need of conservation [6,9,11,12]. Yet, Indications exist that strong conservation measures are not in place to preserve these species and populations continue to decrease in both countries [6-8].

Biodiversity conservation is a complex task which involves different interests of various actors, therefore, it is crucial that all types of stakeholder organizations are involved and interact at different stages of the process [15]. Exchange of scientific information, knowledge and managing experiences among the stakeholder organizations is an important part of these interactions and determine the positive outcomes for biodiversity conservation [16-18]. Social network analysis (SNA) is a commonly used tool to map and quantify these interactions. Social networks, defined as the sets of relationships among the stakeholder organizations, work as channels to facilitate the flow of information and provide opportunities for joint action and collaboration [19-21]. SNA uses a combination of mathematical formulae and models to describe and quantify the existing links among organizations [18]. In recent years, SNA has gained increased attention across a variety of domains including biodiversity conservation [22-24] and proved to be very informative for conservation planning [25].

The social network structure has implications for biodiversity conservation. Social networks can vary in their properties, for example, the number of connections, the structural position of individual stakeholders or the frequency of interactions, and there is no single structure of the network that will be most beneficial in all contexts [26,27]. There are, however, certain network properties which are thought to facilitate effective management of natural resources and effective conservation of biodiversity. For example, a high number of connections in a network was shown to enable improved spread of information relevant to biodiversity conservation [28,29]. Similarly, strong connections, which are frequent connections, are desirable for effective conservation as they indicate high levels of trust [30-33]. Weak connections, which are the less frequent connections, on the other hand, facilitate the transfer of novel information as they tend to connect dissimilar actors [34,35]. Furthermore, the networks in which only one or a limited number of organizations have a central position (holding the majority of relational ties) are more effective for quick mobilization of resources and decision making in the initial phase of conservation action [36,37]. On the other hand, networks with more organizations in a central position are more suitable for long-term environmental planning and complex problem-solving [30]. In summary, whether a network is optimal or not depends on the local context, the organizations involved in the process, and the phase of the conservation process [30,38,39].

Merely the structural analysis of a network (SNA) may not be sufficient to fully understand all the processes and dynamics within the network. Therefore, qualitative analysis of the data provided by the stakeholders is very important to inform and explain the results of the SNA [40]. Qualitative data on the nature and the content of reported interactions as well as the conditions external to the network properties, such as the funding schemes, stability and functioning of organizations, the implementation capacity and the governance arrangements, amongst others provide a deeper understanding of how the network functions and translates into conservation action [39]. Combining the structural analysis of the network data with the qualitative analysis is referred to as the mixed-method approach. The mixed-methods approach has been applied in the context of biodiversity conservation and discussed in more detail by different social scientists [24,41].

Here we employ the mixed-method approach, to study the information sharing network of stakeholders, which are involved in Pontocaspian biodiversity conservation and related information exchange in Romania and compare it to a similar earlier study from Ukraine (S3 Appendix). This study is part of the Horizon 2020 ‘Pontocaspian Biodiversity Rise and Demise’ (PRIDE) program (http://www.pontocaspian.eu/) which was designed to generate scientific knowledge on PC biota and guide effective conservation action. We assess whether the different socio-political contexts in these two countries result in differences in the social network structure of stakeholders, the content of the interactions and the external social variables which may help or hinder the functioning of the network. Importantly, we aim to find how the differences and/or similarities in the two networks translate into PC biodiversity conservation. We conclude the paper with recommendations for improved national and cross boundary conservation efforts. This study contributes to the knowledge on biodiversity conservation and inter-organizational structure of stakeholder institutions in two countries that are politically and culturally different. The combination of quantitative (SNA) and the qualitative analysis provides important additional insights into the network characteristics.

## Materials and Methods

### Stakeholder identification and prioritization

We identified 23 stakeholder institutions in Romania for inclusion in the study, where a stakeholder was defined as an organization who influences or is influenced by the PRIDE research [15]. After engagement, stakeholders which were found to lack any activity or interest in (conservation of) Pontocaspian biodiversity, were omitted, resulting in a final list of 17 institutions (Table 1; Fig 1). We assigned these stakeholders to different categories based on their function and responsibilities: Academic, governmental and non-governmental organizations and a protected area administration (Table 1) following the earlier study from Ukraine (S3 Appendix). The Danube Delta Biosphere Reserve Authority (DDA) is a local administration of the biosphere reserve which was under the Ministry of Environment of Romania by the time of the interview, was transferred and since July 2017 acted under the Romanian Government but is now back to being under the Ministry of Environment. This institution has many functions but most importantly administers the Danube Delta Biosphere Reserve. In the analyses we group the DDA with the other governmental organizations. For Ukraine, we used the dataset compiled earlier consisting of 22 stakeholders of which nine were academic institutions, five governmental organizations, three non-governmental organizations and five protected areas (S3 Appendix).

**Fig 1.**
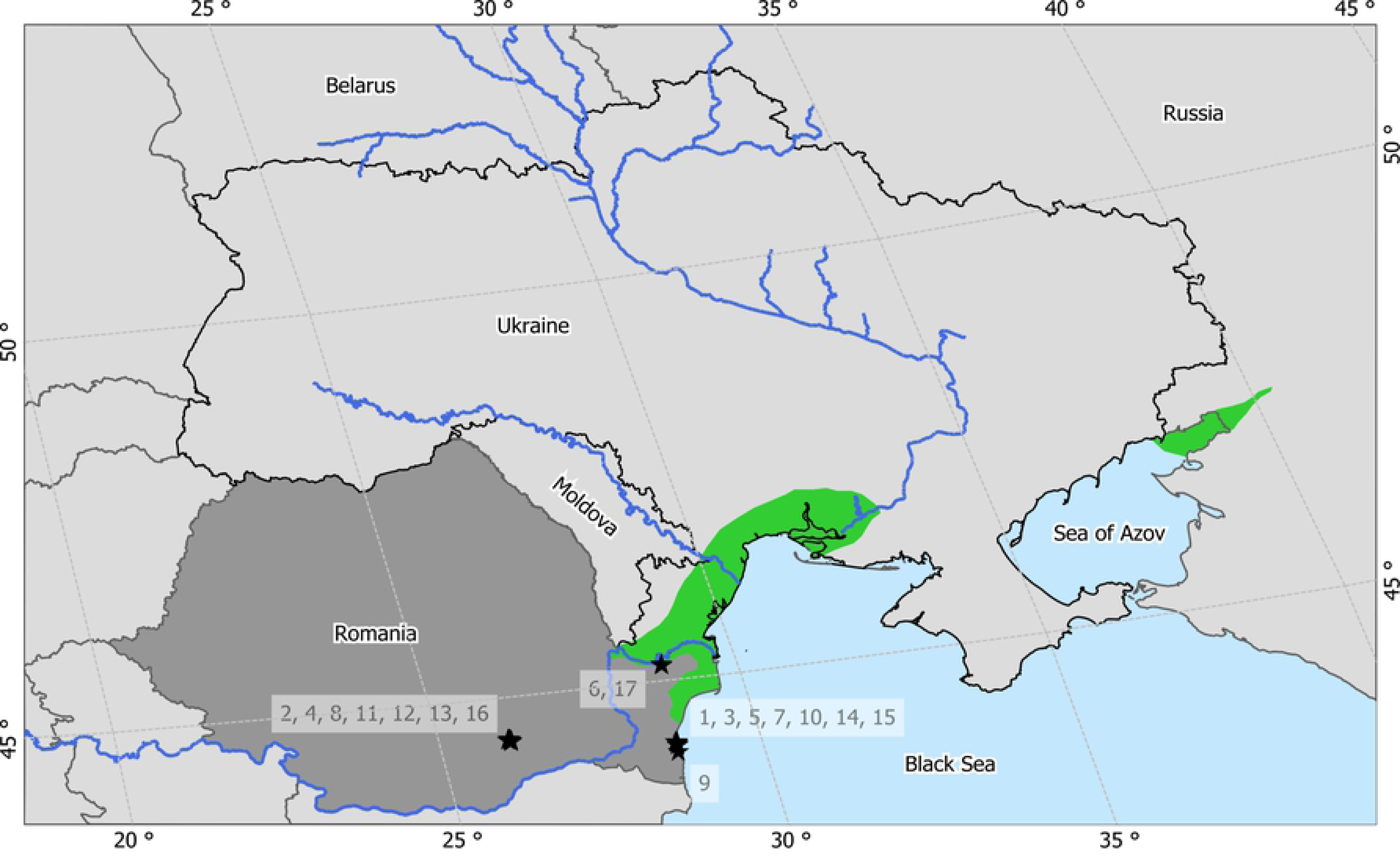
Map of the study area. The black stars on the map represent the stakeholder institutions (IDs in Table 1). Green areas indicate major Pontocaspian habitats.

**Table 1.** List of the 17 selected stakeholders divided into four stakeholder categories.

| ID | Abbreviation | Category | Organization name | Department/Service |
| --- | --- | --- | --- | --- |
| 1 | CMSN | Acad | CMSN Constanța | Delfinariu, Constanta |
| 2 | GAM | Acad | Grigore Antipa National Museum of Natural History |  |
| 3 | GEcM | Acad | GeoEcoMar Constanta |  |
| 4 | IBB | Acad | Institute of Biology Bucharest, Romanian Academy | Department of Microbiology |
| 5 | OUC | Acad | Ovidius University of Constanta | The Faculty of Natural and Agricultural Sciences |
| 6 | DDNI | Acad | The Danube Delta National Institute for Research and Development | Biodiversity Conservation and Sustainable use of Natural Resources |
| 7 | NIMR | Acad | The National Institute of Marine Research and Development "Grigore Antipa" |  |
| 8 | UB | Acad | University of Bucharest | Department of Paleontology |
| 9 | AZS | Acad | Marine Biological Station of Agigea |  |
| 10 | LAC | Gov | Local Environmental Protection Agency in Constanta |  |
| 11 | ANPA | Gov | Ministry of Agriculture and Rural Development of Romania | National Agency for Fisheries and Aquaculture |
| 12 | MOE | Gov | Ministry of environment of Romania | Biodiversity Directorate |
| 13 | MWF | Gov | Ministry of Waters and Forests | Department for Water, Forests and Fishery |
| 14 | MN | NGO | ONG Mare Nostrum |  |
| 15 | OC | NGO | SEOPMM Oceanic Club |  |
| 16 | WWF | NGO | WWF in Romania |  |
| 17 | DDA | Pa/Gov | Danube Delta Biosphere Reserve Authority |  |

Acad represents academic institutions; Gov, governmental organizations; NGO, non-governmental organizations; Pa, protected areas.

### Data collection

We obtained the social network and qualitative data using a survey questionnaire (S1 Appendix). We interviewed the staff members of the institutions or relevant departments during July 2017. Each stakeholder organization was interviewed about each other organization from the list using the same questions.

Qualitatively, we compiled narratives on the context and the content of existing professional links among the stakeholders (both, general and Pontocaspian biodiversity related) and the perceived sufficiency of these links. We extracted the meaning and the content of the interactions from the interviews and no prior data was used. Quantitatively, we collected data on the frequency of those links which were related to PC biodiversity (as defined by the interviewed stakeholders). We used the frequency of contact as a measure of strength (weight) of the relationship (see [37] [42]). We defined five weight categories ranging from no contact to very frequent contact (0-4) and integrated the definitions in the questionnaire as a table (S1 Appendix) so that the interviewees could use it for ranking the strength of the interactions. Answers to the question allowed the generation of a weighted, directed, information and knowledge transfer network.

### Analysis

#### Qualitative analysis

We used the ‘inductive approach’ for qualitative analysis, so the themes (recurrent unifying concepts or statements about the content/subject of the inquiry) were determined based on the collected data and not the prior knowledge or assumptions [43,44]. The themes were established from the collected interviews based on repetitions [45]. We used a ‘constant comparison’ method to refine the dimensions of established themes and to identify the new themes [46]. We then counted the identified themes and determined their relative importance based on the order of frequency.

#### Social network analysis

We translated the collected interviews into an adjacency matrix, a square matrix reporting weights (strength) of all the relational ties. We considered only the confirmed information sharing links - relational links described similarly by both stakeholders involved. The unconfirmed links (14% of all the reported relationships) were considered unreliable and omitted from the study. The values of tie-strengths of confirmed relationships between pairs of stakeholders did not always match. In the case of bi-directional information exchange, tie values were left as reported by the stakeholders. In the case of unidirectional information transfer we selected the lowest and therefore conservative tie value. We imputed the missing network data using the imputation-by-reconstruction method [47]. The preconditions for employing this method are: 1) respondents shall be similar to non-respondents, and 2) the obtained description of the relational link (from the respondent) shall be reliable. A Chi-squared test revealed no significant differences in the distribution of weights of received relationships between the respondents and non-respondents (*p*-value = 0.92), meaning that respondents are similar to non-respondents. Furthermore, the confirmation rate (proportion of relational links described similarly by both nodes involved) was 85 % indicating that the descriptions of relational links (provided by the respondents) can be considered as reliable. Therefore, we used the reconstruction method to impute the missing ties in the network. We visualized the sociogram using the CRAN R package ‘igraph’ [48].

We calculated the basic network characteristics such as number of actors and relational ties, graph density and centralization using CRAN R package ‘igraph’ [48]. The mean shortest distance was calculated using the CRAN R package ‘tnet’ [49] because the ‘igraph’ package does not take edge weights into account when measuring the shortest distance. Graph density is the extent to which nodes are connected to each other in the network. It is calculated by dividing the number of existing ties by all the possible ties in a network [18,50]. Network centralization is the extent to which certain actors are more connected in the network than the others [18,51]. A centralized network is one where only one or few actors are having the majority of the ties. Such a network has a high overall centralization score (on a 0 to 1 scale, 0 being completely decentralized and 1 fully centralized). Shortest distance is a minimum number of steps that the nodes are away from each other in a network; in weighted networks the tie weights are taken under consideration [52]. We used frequency of contact as a measure of strength of the relationship and defined strong relationships as the weights higher or equal to 3 on a scale ranging from no contact to very frequent contact (S1 Appendix).

We measured the centrality of individual nodes using degree centrality and betweenness centrality values. Degree centrality is the number of connections a particular actor has with all the other actors in a network [53]. We calculated the degree of a node through an in-degree and out-degree values. In-degree of a node is the number of in-coming links to it from the other nodes in a network and the out-degree of a node is the number of out-going links from this node to the other nodes in a network [54]. Furthermore, we measured and used the node strength values (extension of the degree centrality to the sum of tie weights when analyzing weighted networks) to determine the size of the nodes in a sociogram [33,55,56]. Betweenness centrality measures the extent to which a node is among other nodes in a network [53]. For weighted networks the betweenness centrality measure is based on algorithm of shortest path distance [57,58] which was lately further developed to integrate the cost of intermediary nodes in the formulae [52]. We calculated node-level statistics using the CRAN R package ‘tnet’ [49] which considers tie weights and corrects for the number of intermediary nodes. We regarded the central stakeholders as the ones with centrality scores higher than the third quartile threshold values [23,42,59].

We measured brokerage combining quantitative and qualitative approaches. Brokers are the nodes which are between other nodes in a network and have the power to control the flow of information [34,60,61]. Quantitatively, brokerage was measured through the betweenness centrality and the Burt’s constraint metric [34,60]. Betweenness centrality locates the brokers structurally, with respect to all the other actors in the network. Burt’s constraint, however, is a local measure of brokerage based on the triadic closure principle. A node connecting two disconnected nodes in an incomplete triad has a power to broker. Such nodes have low Burt’s constraint score, i.e. their behavior is not constrained by the other disconnected nodes in a triad [17,61]. Qualitatively, we examined the network narratives and searched for the evidence that the stakeholders are actually engaging in brokering behavior. Brokering behavior in the context of biodiversity conservation implies the mobilization of information, deliberation between different types of stakeholders and potentially the mediation through working groups to address conservation issues [62]. In our study, we regarded the stakeholders with high betweenness scores, which also accounted for low Burt’s constraint values, and were involved in brokering behavior as brokers. We used only the strong ties (≥ 3) to calculate betweenness centrality and Burt’s constraint metric as they reflect regular contacts. We calculated Burt’s constraint utilizing CRAN R package ‘igraph’ [48].

A null-model test was used to identify the presence of ‘network homophily’ in the network. ‘Network homophily’ is the selective linking between actors based on specific attributes, in our case the category of stakeholder institutes [63]. With a null-model test, we tested whether densities within and between stakeholder groups (defined by the stakeholder category) were significantly higher or lower than the random expectation. We randomly assigned nodes to the stakeholders proportional to the true network and subsequently assessed the stakeholder’s within and between group densities replicated 1000 times, resulting in 1000 stakeholder group density values. We ranked the obtained 1000 random values from low to high and compared the actual within and between group densities to the randomized results. If the actual density values were larger than the upper or smaller than the lower 2.5% threshold value of the random distribution, we regarded the true within or between group densities to be significantly higher or lower than expected by random chance.

#### Ethics statement

The social network analysis of stakeholder organizations which we conduct here is not subject to ethical screening as it is, for example, for medical and/or socio-medical studies, which provide personal data. As such, we did not seek a priori ethics review nor is there any established procedure within our organization (Naturalis Biodiversity Center) which we could follow. We informed all participants prior to the interviews that they were being interviewed on behalf of the organization which they were affiliated to, and that the results would be part of a publication, assuring them that no participant would be individually identifiable and asked them whether they objected. All the interviewed stakeholders understood and did not object to analyses and publication of their responses.

## Results

Here we report on the results from the conservation network from Romania, which we will compare to the Ukrainian network (S3 Appendix). Out of the 17 Romanian institutions 15 were interviewed (covering 88% of the network data): 14 through face to face in-depth interviews and 1 through an electronic questionnaire via email. From the missing two institutions one was met, but interviewing was not possible (Table 1). The remaining institution could not be reached.

### Network functionality and content of interactions

The studied network in Romania interacts on multiple levels on a variety of topics (S2 Appendix). Unlike in Ukraine, the majority of interactions in Romania were based on projects. Some of the joint projects were described as “the EU funded projects” by the stakeholders (S2 Appendix). In other cases, the source of financing was not specified. Outside the projects, the exchange of comprehensive data in Romania was subject to payment, because detailed information was not freely available. In some cases, relationships were reported to be unclear due to recent changes in institutional arrangements and governance. Exchange of data involving the payment and unclear relationships due to institutional rearrangements were also reported to be reasons for insufficient collaboration (discussed in detail below under the ‘Perceived sufficiency of collaboration’).

In Romania, like in Ukraine, Pontocaspian species played a role in just a few of the inter-organizational relations (S2 Appendix). The exchange of information with regard to PC biodiversity was mostly project-based in Romania. These projects, however, mostly focused on the flagship species such as sturgeons (e.g. ‘LIFE for Danube Sturgeons’ project). Furthermore, monitoring according to the EU Habitats Directive (Article 17), includes the PC habitats and sturgeon species. Outside the projects the exchange of information mostly occurred through joint scientific work (fieldwork, publications) or was subject to payment for data. The majority of the relationships related to PC biodiversity involved PC habitats or species as a minor component of the interaction.

### Network structure

The Stakeholder network in Romania was not well connected (Table 2; Fig 2) with a total number of 57 relational ties out of 272 potential ties resulting in a network edge density measure of 21% (Table 2). For comparison, the PC network in Ukraine had the edge density value of 41%. On average each organization in Romania had 7 relational ties with other stakeholders in the network, while in Ukraine each stakeholder had 17 ties. This resulted in bigger mean distance between stakeholders in the Romanian network compared to Ukraine (2.5 in Romania vs 1.5 in Ukraine). The Romanian network had a lower degree of centralization score (22%) than the Ukrainian network (38%). However, based on individual node statistics (presented below) we define both networks to be centralized networks. The correlation of incoming and outgoing ties, although positive in both networks, was lower in Romania compared to Ukraine (rho = 0.31 in Romania vs. rho = 0.78 in Ukraine) indicating that information exchange is in general less reciprocated in Romania (Table 2). When governmental organizations (including the DDA) were omitted from the Romanian network, the correlation increased (rho = 0.79), suggesting that the governmental organizations received information from multiple sources but did not share similarly. In Ukraine, the exclusion of governmental organizations from the analysis did not make a big difference (rho = 0.76 after exclusion) suggesting that governmental organizations in Ukraine were more open to sharing information. The majority of all the relationships were strong (weights 3 or 4) in both countries (56% in Romania and 61% in Ukraine).

**Fig 2.**
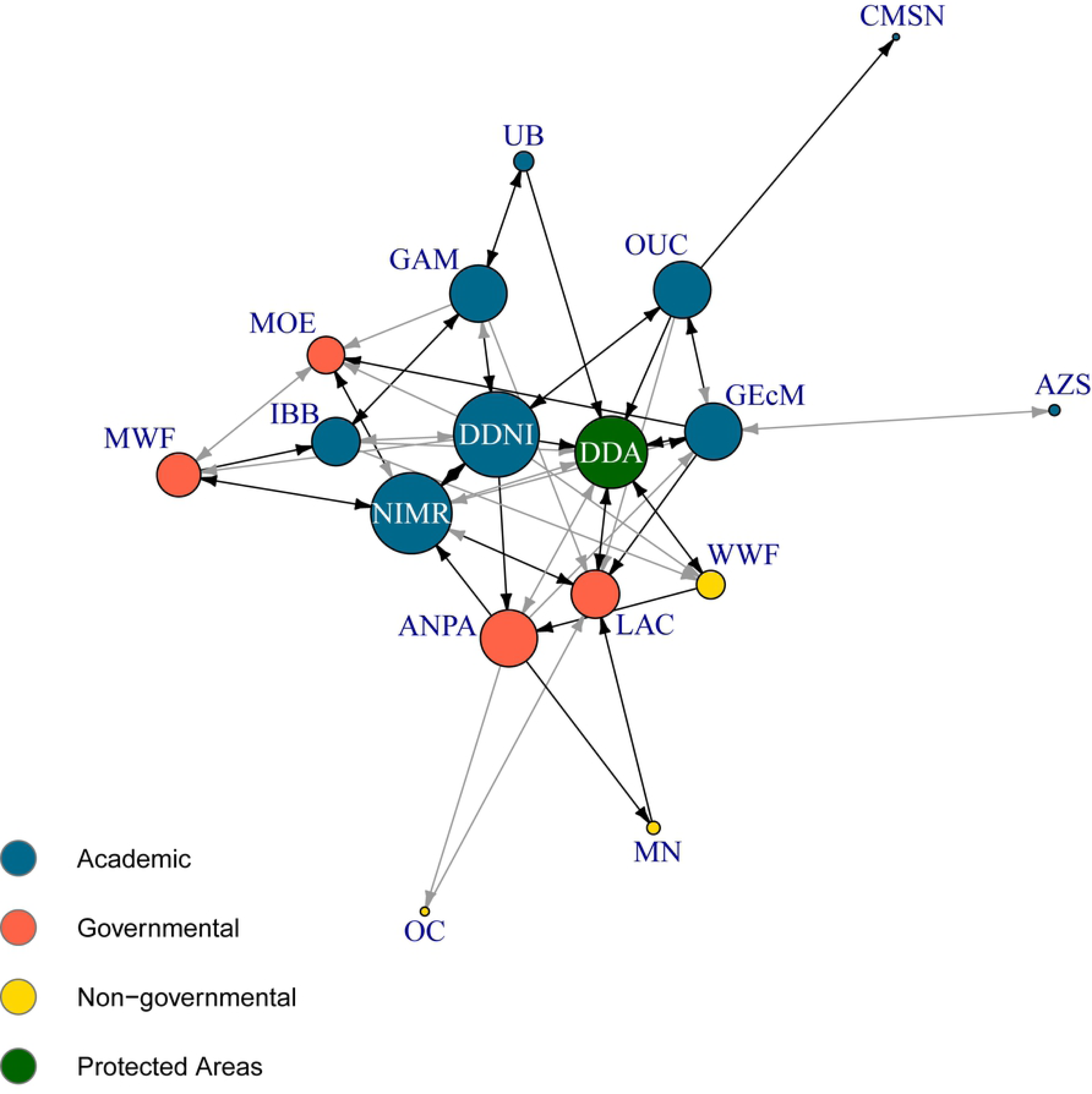
Sociogram of Romanian stakeholders involved in Pontocaspian biodiversity conservation and conservation planning. Nodes represent organizations (see Table 1 for institution descriptions). The size of the nodes corresponds to the node strength (sum of weights of all its links). Arrows represent relationships between the nodes. The black arrows, ties with values ≥3, represent strong relationships. The gray arrows, ties with value < 3, represent weak relationships.

**Table 2.** Network statistics for Romanian stakeholder network compared to the Ukrainian stakeholder network (S3 Appendix).

| Network data | Romania | Ukraine |
| --- | --- | --- |
| Total actors | 17 | 22 |
| Total No. of ties | 57 | 191 |
| Mean degree | 7 | 17 |
| Density (%) | 21 | 41 |
| Degree of centralization (%) | 22 | 38 |
| Tie reciprocity (rho) | 0.31 | 0.78 |
| Tie reciprocity (rho) excluding the Gov. organizations | 0.79 | 0.76 |
| Strong/weak ties (%) | 56/44 | 61/39 |
| Mean shortest distance | 2.5 | 1.5 |

### Central stakeholders

We found five central stakeholders in Romania and six in Ukraine, based on degree centrality scores (Table 3). In both networks three academic institutions out of nine had a degree centrality score higher than or equal to the third quartile threshold value (≥9 in Romania and ≥20 in Ukraine), indicating high involvement of these organizations in the exchange of relevant information. Unlike in Ukraine, where the major decision-making organization (Ministry of Ecology) was the most central stakeholder, in Romania, the analogous institution (Ministry of Environment) was not actively involved in exchange of relevant information. Instead, the Local Environmental Protection Agency in Constanta (LAC) was the central governmental institution with high degree centrality score. The Danube Delta Biosphere Reserve Authority (DDA) in Romania and the Danube Biosphere Reserve Administration (DBR) in Ukraine were both active in the network with high degree centrality scores. The non-governmental organizations were marginally involved and had few connections in both countries. All the central stakeholders in Ukraine had more strong than weak connections. In Romania this was also the case with an exception of GEcM which had equal amount of weak and strong ties.

**Table 3.**
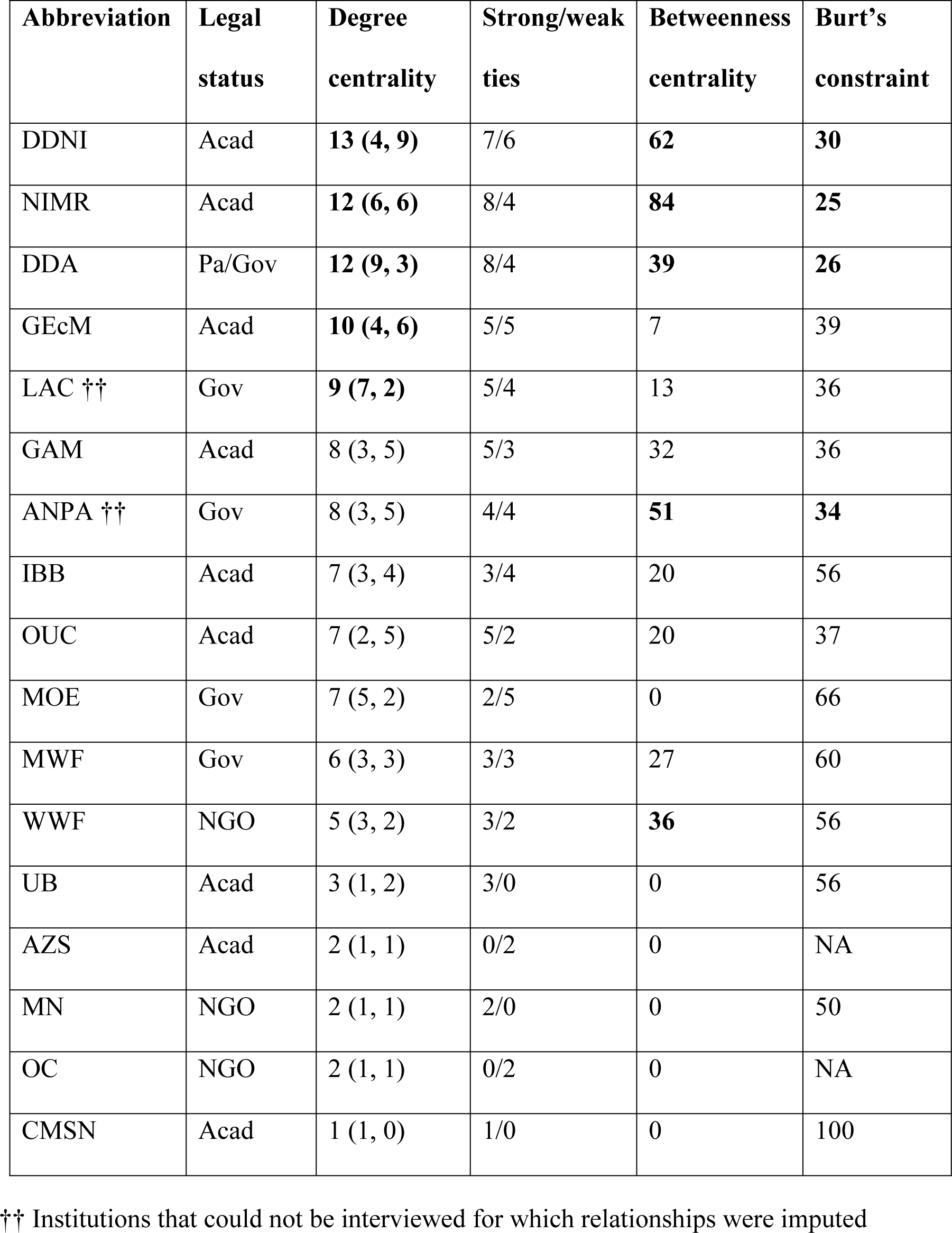
Node-specific measures for the network. The values in the brackets under the ‘Degree centrality’ represent the in-degree and out-degree measures respectively. In bold are the values higher than, or equal to the third quartile threshold (lower or equal to the first quartile threshold in case of ‘Burt’s constraint’). Burt’s constraint values for AZS and OC are not defined (NA) as the calculation was based only on strong ties (≥ 3).

| **Abbreviation** | **Legal status** | **Degree centrality** | **Strong/weak ties** | **Betweenness centrality** | **Burt's constraint** |
| --- | --- | --- | --- | --- | --- |
| DDNI | Acad | **13 (4, 9)** | 7/6 | **62** | **30** |
| NIMR | Acad | **12 (6, 6)** | 8/4 | **84** | **25** |
| DDA | Pa/Gov | **12 (9, 3)** | 8/4 | **39** | **26** |
| GEcM | Acad | **10 (4, 6)** | 5/5 | 7 | 39 |
| LAC †† | Gov | **9 (7, 2)** | 5/4 | 13 | 36 |
| GAM | Acad | 8 (3, 5) | 5/3 | 32 | 36 |
| ANPA †† | Gov | 8 (3, 5) | 4/4 | **51** | **34** |
| IBB | Acad | 7 (3, 4) | 3/4 | 20 | 56 |
| OUC | Acad | 7 (2, 5) | 5/2 | 20 | 37 |
| MOE | Gov | 7 (5, 2) | 2/5 | 0 | 66 |
| MWF | Gov | 6 (3, 3) | 3/3 | 27 | 60 |
| WWF | NGO | 5 (3, 2) | 3/2 | **36** | 56 |
| UB | Acad | 3 (1, 2) | 3/0 | 0 | 56 |
| AZS | Acad | 2 (1, 1) | 0/2 | 0 | NA |
| MN | NGO | 2 (1, 1) | 2/0 | 0 | 50 |
| OC | NGO | 2 (1, 1) | 0/2 | 0 | NA |
| CMSN | Acad | 1 (1, 0) | 1/0 | 0 | 100 |

†† Institutions that could not be interviewed for which relationships were imputed

Four of the six central stakeholders in Romania (DDNI, NIMR, DDA and ANPA) had a structurally favorable position to act as brokers based on both the betweenness centrality and the Burt’s constraint scores (Table 3). Qualitative data, however, showed that these structurally well-positioned organizations were not engaging in brokering behavior with regard to Pontocaspian biodiversity (see below). In Ukraine, on the other hand, two out of four structurally well positioned organizations were found to actually engage in brokering behaviors. The major decision-making organization (Ministry of Ecology) was the biggest broker of the Ukrainian network, followed by an academic institution (Institute of Marine Biology). The Ministry of Ecology in Ukraine was found to have various working groups relevant to PC biodiversity conservation, to actively communicate and exchange information and bring otherwise disconnected stakeholders together for conservation planning. Similarly, the Ukrainian Institute of Marine Biology (IMB) which is a scientific curator for several protected areas coordinated their research and connected them, which were otherwise disconnected or very weakly connected (S3 Appendix).

WWF accounted for high betweenness values in both networks; however, they did not directly bridge many disconnected nodes (indicated by their high Burt’s constraint scores). The qualitative data showed that WWF Romania and WWF in Ukraine were actively involved in the conservation of sturgeon species through the enforcement of conservation laws and awareness raising. They had large number of volunteers in both countries and sometimes brought the otherwise disconnected stakeholder organizations together for joint conservation action. Their work, however mostly focused on charismatic PC species and the wider PC taxa was absent from their conservation agenda.

### Network homophily

Across the Romanian network, different stakeholder categories had various tie densities, but connectedness was not significantly higher than expected from the null-model test, indicating the absence of network homophily (Table 4). In Ukraine, we found strongly connected academic institutions with significantly higher within group density value than expected by chance suggesting network homophily. The non-governmental organizations were marginally involved in the exchange of Pontocaspian information in both, Romanian and Ukrainian networks. In Romania, NGOs were significantly less connected to the academic institutions than expected by chance and had no links among themselves. In Ukraine, NGOs were also significantly less connected to academic organizations and had only two links among themselves (Table 4). The density values within and between other groups of stakeholders were not significantly different from random expectation. The academic organizations had more strong than weak connections among themselves in both networks indicating regular exchange of information within this group. Furthermore, they were strongly connected with the governmental organizations in Romania, but less so in Ukraine. Governmental organizations were more strongly connected to each other in Ukraine than in Romania, and strongly connected with the NGOs in both countries. Most of the very few connections NGOs had with each other and with academia were strong in Ukraine, unlike in Romania.

**Table 4.**
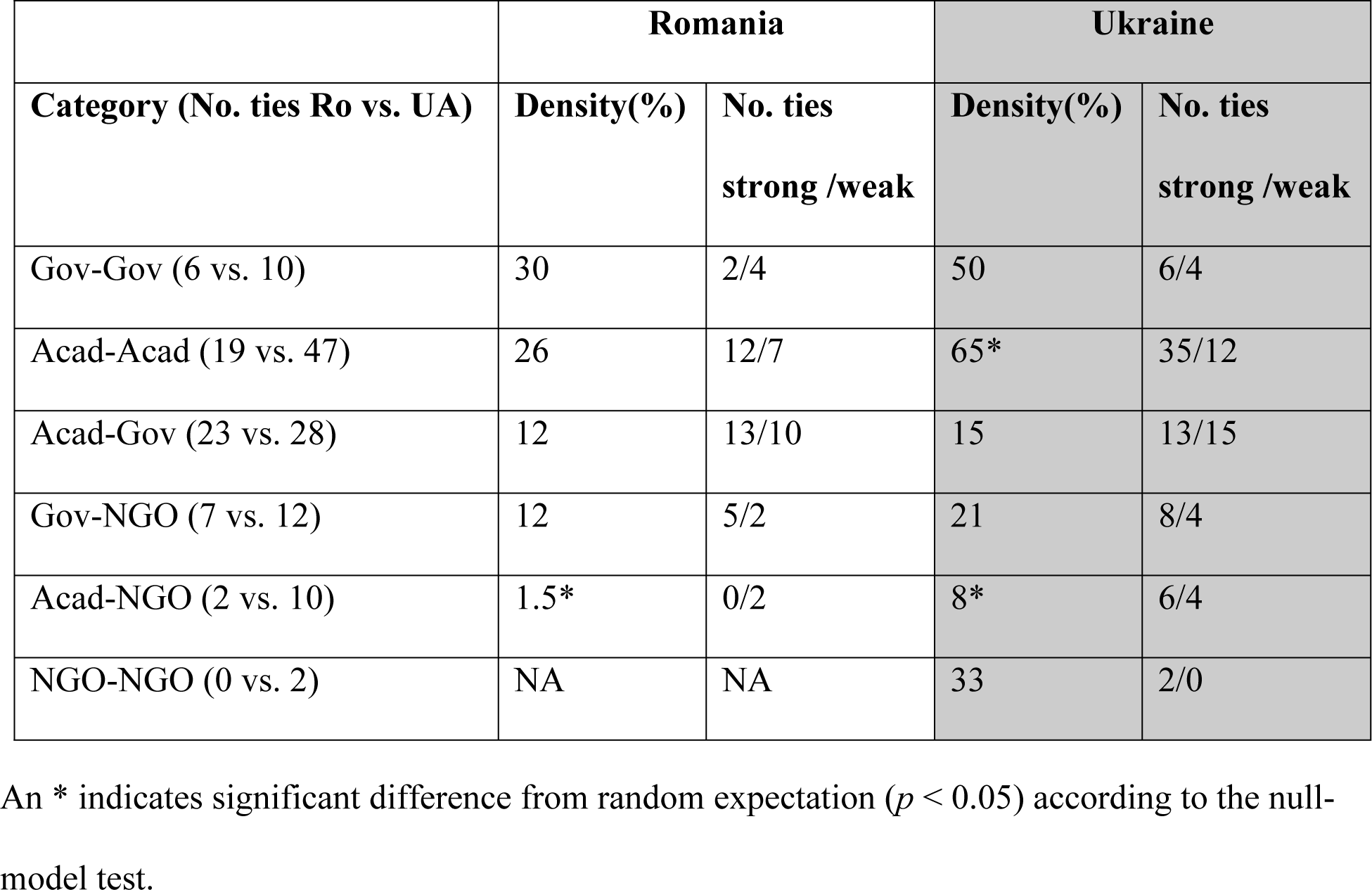
Density of the ties within and between stakeholder categories in Romania and Ukraine. The values in brackets under the ‘Category (No. ties RO vs. UA)’ represent the number of existing relational ties in Romania and in Ukraine.

### Perceived sufficiency of collaboration

In both networks, the majority of the relationships among the interviewed stakeholders were described to be sufficient to exchange information and to achieve effective collaboration (54% in Romania vs. 68% in Ukraine), the remainder to be insufficient (Table 5). Some of the reasons for insufficient relationships were similar in both countries. For example, ‘budget constraints’ was the most prominent factor in both countries limiting more intense collaboration (26% in Romania vs. 45% in Ukraine). ‘Lack of interconnection’ was another factor reported in both countries (16% in Romania vs. 8% in Ukraine) to complicate collaboration. ‘Lack of interconnection’ referred to the situation in which one party is interested and eager to have more collaboration/exchange of information, while the other party is not responding due to, for example, different interests or priorities. Other reasons given for insufficient collaboration were different in two countries. Importantly, free exchange of data was common in Ukraine, but was indicated as a limiting factor in Romania (20%) as it was not freely available. Similarly, ‘political constraints’ (18%), ‘institutional turnover’ (14%) and ‘institutional competition’ (6%) were reported only in Romania as factors hampering the establishment of relationships and collaboration. ‘Political constraints’ mostly referred to governmental organizations being influenced by politics, and being closed for consultations with the academic sector or non-governmental organizations. The ‘legal limitations’ which in Ukraine mostly referred to the contradicting national laws (S3 Appendix) were not mentioned in Romania.

**Table 5:** Perceived sufficiency of reported professional interactions among stakeholder organizations in Romania and Ukraine. The sub-themes in italics are under the theme ‘Insufficient’.

| **Perceived relationships** | **Romania** | **Ukraine** |
| --- | --- | --- |
| Sufficient (%) | 54 | 68 |
| Insufficient (%) | 46 | 32 |
| *Budget constraints (%)* | 26 | 45 |
| *Limited exchange of information (%)* | 20 | NA |
| *Political constraints (%)* | 18 | NA |
| *Lack of interconnection (%)* | 16 | 8 |
| *Institutional turnover (%)* | 14 | NA |
| *Institutional competition (%)* | 6 | NA |
| *Legal limitations (%)* | NA | 29 |
| *Unknown (%)* | NA | 13 |
| *Employee turnover* | NA | 5 |

## Discussion

Conservation of Pontocaspian (PC) biodiversity is critically dependent on adequacy of conservation measures and the coordination of actions across their distribution range - the Northern part of the Black Sea and the Caspian Sea region. This paper aims to assess the adequacy of the stakeholder networks for conservation in two countries responsible for a large part of the native range of PC biota. We compare the social network structures of stakeholders involved in biodiversity conservation in Romania and Ukraine, based on new data from the former and the older data from the latter (S3 Appendix). First we discuss the implications of the new, Romanian results for effective conservation and compare it then to Ukraine. We examine the challenges within, as well as beyond the network structure for optimal PC biodiversity conservation and provide recommendations for improved cross border conservation efforts.

### Implications of the Romanian network properties

Both the interviewees and the literature [6,7] indicate that Pontocaspian biodiversity is declining in Romania and conservation measures are not always in place, suggesting that the studied network translates into sub-optimal conservation action [6,7,10]. The structure of the conservation network may be one of the underlying factors (Table 2; Fig 2). The Romanian network properties such as few connections, low tie reciprocity, and unrealized potential of central stakeholders, suggest suboptimal PC biodiversity conservation conditions. Conservation action is further challenged by external social variables such as instability of the institutional arrangements and governance as well as funding limitations and political constraints (Table 5). Interestingly, stakeholder interactions rarely target conservation of PC biodiversity directly (S2 Appendix). This indicates a low priority for the conservation of PC biota. Many links relevant to biodiversity conservation in general may benefit PC species indirectly, while some links target PC flagship species such as sturgeons. For example, biodiversity monitoring activities (according to the EU Habitat Directive) include PC habitats and the “LIFE for Danube Sturgeons” project is specifically targeting sturgeon conservation.

The Romanian network is centralized, with few stakeholders across different stakeholder categories holding the key positions (Tables 2 and 3; Fig 2). Whether a centralized network is suitable for effective conservation action depends on the phase of the conservation process [30,38]. Decentralized networks are in general suitable for long-term environmental planning and complex problem solving, due to the need of multiple stakeholders (across the disciplines) contributing to the solution of a problem, providing different knowledge and perspectives [30]. A more centralized network (with one or few very central stakeholders) can be effective in the initial phase of the conservation process when resources need mobilization and the coordination of joint action is required. In Romania (and also in the Ukraine) research on Pontocaspian biodiversity has a long history [6,9], but the translation of research outputs to conservation action is relatively novel and not always in place [6,9,64]. We suggest that in the current phase of PC biodiversity conservation in Romania a centralized network is likely an optimal network. Although, it is instrumental that the central actors use their abilities and structurally favorable positions in practice to take a coordinating role in resolving conservation challenges. Additionally, more involvement of an influential decision-making organization, such as the Ministry of Environment, would be desirable for the network to effectively address conservation action. In general, the central organizations in Romanian network do not exploit their positions to initiate collective action. For example, while DDNI, NIMR and DDA are potentially the brokers in the network (Table 3), we found no evidence, of any action related to brokerage. This may be due to lack of appropriate incentives or the limited knowledge on the need for conservation of PC biodiversity.

Not all stakeholder categories are well embedded in the structure of the Romanian conservation network. For example, the NGOs are marginally involved in the network (Table 4) and governmental organizations have limited numbers of reciprocated ties (Table 2). This may suggest low motivation of governmental and non-governmental stakeholders to engage in Pontocaspian conservation action [65]. The marginal involvement of NGOs has been observed in earlier study in Romania in the Natura 2000 governance network [66]. The lack of reciprocated communication (governmental stakeholders receiving information from multiple sources but not sharing back to the network) may be indicative of strong hierarchy [67]. This idea is supported by interviewees, which mention ‘political constraints’ and ‘lack of interconnection’ as factors limiting collaboration (Table 5). Effective biodiversity conservation requires information exchange between diverse stakeholder categories [32,37]. Therefore, more involvement of NGOs and governance actors may benefit conservation of PC biodiversity.

The optimal functioning of the network is further challenged by the external conditions such as funding scarcity, political constraints and institutional turnover (Table 5). Funding for conservation is scarce and limits collaboration. Beyond the few funded projects, information is not often exchanged and data availability is limited. In addition, scientific organizations that hold most biodiversity information (e.g. DDA, DDNI and GAM), are insufficiently funded by the government and data quality, availability and persistence is dependent on their success to find additional funding. Limited funding discourages collaboration and increases competition. For example, several stakeholders involved in research and conservation have their own research vessels (e.g. NIMR, GEcM, OC) which are expensive to maintain, yet they do not coordinate their activities because they are competing for funds. Furthermore, the institutional alignment of stakeholder organizations is not stable in Romania, complicating the maintenance of relations and building trust. For example, the Ministry of Environment (MOE) and Ministry of Waters used to be one, but were recently split up; the DDA was transferred from the MOE to central government by the time of the interview but is now back to being under the MOE; and the Marine Biological Station of Agigea became independent after being a research station of the University of Iasi. These changes complicate the formation and maintenance of relationships. Continuous institutional reforms to adjust to the EU structures has been reported to not always have positive outcomes in Romania [68]. Finally, ‘political constraints’ and ‘lack of interconnection’ exacerbate the lack of exchange of information in Romania.

### Comparing Romania and Ukraine

The Romanian network properties can be compared with those of Ukraine (S3 Appendix) as the data collection has followed identical approaches. The Romanian network is slightly smaller (we identified 17 stakeholder organizations compared to 22 in Ukraine). Importantly, the surface area of Pontocaspian habitats in Ukraine is much larger than in Romania. PC species are in need of conservation, however, targeted conservation is limited in both countries and seem suboptimal. Conservation actions in Romania are mostly project based (and often EU-funded), unlike in Ukraine. While information exchange in Ukraine is common outside the formal projects, this is not the case in Romania. In Ukraine, academic institutions (under the NASU), are not allowed by law to sell data and are obliged to share information for free. Furthermore, all the protected areas and academic organizations are obliged to share the biodiversity related information with the Ministry of Ecology (MENR) – sometimes directly (through specific requests) or indirectly (through different departments of Regional Administrations). The MENR and other governmental organizations, in turn, are open to communicating the information back to the network (S3 Appendix). This results in much more free exchange of information and data and in a higher number of reciprocated connections in Ukraine compared to Romania. Data and information sharing are preconditions for effective conservation action, and the Ukrainian network seems to be better set-up to achieve it.

The network properties and reported interactions are very different in two countries. In Ukraine, the network is structurally strong with many connections, highly reciprocated ties and clearly defined broker institutions (S3 Appendix). In Romania, the network is structurally weaker. The content of interactions is not favorable in either of the countries for effective Pontocaspian biodiversity conservation, as this biota is not a direct target of interactions. Both Ukrainian and Romanian networks are centralized; the difference is that in Ukraine the major decision-making organization (Ministry of Ecology) is also in the most central position. The analogous organization from Romania (MOE) is not one of the central stakeholders. This may suggest that on the governmental level PC biodiversity conservation has a lower priority in Romania, since the Ministry is the major decision making and coordinating organization for biodiversity conservation and conservation planning in both countries. Additionally, the Ministry of Environment and other governmental organizations in Romania are less involved in consistent, bi-directional exchange of information (Table 2) compared to Ukraine indicating more openness of Ukrainian governmental organizations to collaboration. Similarly, the central stakeholders are not exploiting their favorable positions in Romania as much as they do in Ukraine, suggesting lack of incentive for Romanian stakeholders to initiate PC biodiversity related conservation action, which, in turn may be due to the project-based interactions in Romania and limited funding availability for non-charismatic PC species. Funding scarcity for the non-charismatic PC species can also explain the marginal involvement of NGOs in both countries.

In both countries, there are limited funding options available for biodiversity conservation and the sources of funding are diverse. The main difference is that in Romania collaboration declines when a project is over, while in Ukraine organizations continue to collaborate and exchange information beyond projects. This may be partly because they have legal obligations to do so. From network narratives, we learned that Romanian stakeholders are involved in many more projects than Ukrainian stakeholders, and many of these projects are EU funded. Yet, the Romanian network is less dense than the Ukrainian one. This finding may indicate that the network in Romania is more reactive rather than proactive.

Legal limitations in Ukraine are complicating collaboration and, in some cases, result in unfavorable institutional alignments which is not the case in Romania. Legal limitations in Ukraine refer to the contradicting laws creating confusion in research methodology and collaboration frameworks. It also refers to uncoordinated action of regional administrations (i.e. not in line with the Ministry of Ecology) such as issuing permissions and acting without consulting the Ministry (S3 Appendix). The legal limitations were not reported to be complicating collaboration and exchange of information in Romania, which may suggest more consistency in the policies in Romania, which in turn may be resulting from the processes of harmonization to the EU Acquis. In Ukraine, refining the national legislation and approximation to the EU Directives were mentioned several times by the stakeholders as they narrated about the content of their interactions. Therefore, improvements can be expected in the coming years in Ukraine with regard to the national legislation which are likely to result in more coordinated action among stakeholder organizations. Finally, while the institutional reforms are common in Romania, the Ukrainian network is very stable (Table 5). Specific reasons underlying the institutional turnover in Romania were not mentioned by the interviewed stakeholders but the fact itself was reported to be a challenge for establishing relationships and conducting consistent work.

In summary, the road to optimal Pontocaspian biodiversity conservation is different in Ukraine and Romania. In Ukraine, the network is structurally close to optimal, which is a necessary base for effective conservation, but the conservation is mostly challenged because of the untargeted approach and external social variables such as funding scarcity and inconsistent policy frameworks. These external challenges are not specific to PC biodiversity, but concern the entire biodiversity in Ukraine. In Romania, there is a room for improvement within the structure of the network as well as beyond. Within the network, more involvement of governmental organizations in the coordinating roles, and engagement of central, information holding organizations in the brokering behavior could result in a stronger network with a higher potential for optimal conservation action. Beyond the network, funding scarcity, political constraints, institutional turnover and difficulty of detailed information exchange are the obstacles. The common challenge in both countries is that PC biodiversity conservation has a low priority and awareness raising is necessary.

### Coordinating joint Pontocaspian biodiversity conservation actions

Romania and Ukraine share the Danube Delta, the Black Sea coastline and associated habitats in which Pontocaspian biota occurs which may benefit from a coordinated action of both countries. Some of the PC species, e.g. the sturgeon species, are mobile and not limited to the administrative and political boundaries. Furthermore, PC species have a patchy distribution in Ukraine and Romania and face similar pressures in these two countries. Cross-border collaboration is therefore instrumental to achieve common conservation objectives and optimal conservation action. Sharing the management experiences and best practices among the organizations from both countries can help to the development of common organizational awareness and embolden joint efforts and understanding.

The great significance of cross-border collaboration has been recognized by international conventions and the EU, which resulted in several cross-border cooperation projects [69]. In our interviews we did not specifically address cross-border collaboration between Romania and Ukraine with regard to PC biodiversity, but from the network narratives we learned that institutions in both countries are aware of each other and some collaboration exists. Main established programs relevant to PC biodiversity conservation are the cross-border cooperation (within the European Neighborhood Instrument - https://www.euneighbours.eu/en) and the LIFE program of the EU. The former includes “Black Sea”, “Danube”, and other bilateral or trilateral (+ Moldova) ecological programs with large budgets. Usually in their formulations the term “Pontocaspian” does not exist, but the projects mainly concern the habitats of the PC fauna (Danube Delta and Prut River, Lower Dniester and the Black Sea coastline of Ukraine, Romania and Bulgaria). The LIFE program targets the Danube sturgeons. For other PC taxa we did not find evidence for deep collaboration. PRIDE was a pioneering EU funded project which attempted to broaden the sturgeon network in Ukraine to include other PC taxa in the awareness raising activities of WWF in Ukraine. It is important that there are more projects in future, which can extend the current Pontocaspian networks in Ukraine and Romania to the entire PC biota. Such projects can be expected to raise awareness and increase the interest of governmental and non-governmental organizations to collaborate more and exchange the relevant information.

## Conclusions

We conducted this study as part of the PRIDE project to examine the current inter-organizational structure of stakeholders in Romania and understand the implications of network characteristics for the threatened Pontocaspian biodiversity conservation. We compare the results from Romania to an earlier study from Ukraine as these two countries share the responsibilities to conserve Pontocaspian habitats and species but their legal and political frameworks are different. We found that the social networks of stakeholder organizations in Romania and Ukraine are very different - both, the structure as well as the content and the context of interactions differ. Structurally, Ukrainian network is strong, whereas in the Romanian network there is a room for improvement, through e.g. more involvement of governmental and non-governmental organizations and increased motivation of central stakeholders to initiate conservation action. Regardless, both networks translate into sub-optimal Pontocaspian biodiversity conservation action and the road to effective conservation is different in two countries. In an earlier study in Ukraine, we concluded that the maintenance of existing network is a necessary base, and can be expected to result in optimal conservation action if the content of interactions (through awareness raising and capacity building) and external social variables (funding and legal limitations) are improved. In Romania, the external social variables (institutional turnover, political constraints and funding scarcity) have a higher influence on the network structure than in Ukraine resulting in complicated data exchange, fewer connections and a hierarchical governance system. The current network structure therefore cannot be expected to be effective in addressing the Pontocaspian biodiversity conservation without the involvement of governmental and central stakeholder organizations in coordinating roles. Fostering the cross-border collaboration through new calls for project proposals, which involve wider Pontocaspian taxa, will likely increase the awareness and interest of different types of stakeholders to engage more in the conservation action related to this biota. Extending the Sturgeon networks to the other, non-charismatic Pontocaspian species may be a preferable course to initiate such action.

## Acknowledgments

We would like to thank Professor Marius Stoica and Lea Rausch from the University of Bucharest, and Dr. Luis Ovidiu Popa, Alberto Martinez Gandara, Oana Paula Popa and Ana-Maria Krapal from the Grigore Antipa National Museum of Natural History for their invaluable assistance and support in Romania and helping to arrange meetings with stakeholders. We also thank all the interviewed stakeholders for finding time to meet and for providing honest and thorough answers to the survey questions. We acknowledge Cristina Sandu from the International Association for Danube Research for helpful comments on earlier drafts of this paper. We would like to also acknowledge our colleagues from Naturalis Biodiversity Center: Nieke Knoben, for the assistance in developing the questionnaire and Caroline van Impelen for organizational support.

## Author contributions

Conceptualization: AG, NR, KB, FPW. Data collection: AG. Formal analysis: AG, NR. Methodology: AG, NR, KB, FPW. Validation: NR, KB, CI, BP, MOS, NG, VA, FPW. Writing – Original Draft: AG, NR, KB, FPW. Writing – Review & Editing: CI, BP, MOS, NG, VA.

